# Intervening to preserve function in ischemic cardiomyopathy with a porous hydrogel and extracellular matrix composite in a rat myocardial infarction model

**DOI:** 10.1101/2024.07.02.601690

**Authors:** Yasunari Hayashi, Taro Fujii, Seungil Kim, Stephen F. Badylak, Antonio D’Amore, William R. Wagner

**Affiliations:** McGowan Institute for Regenerative Medicine, University of Pittsburgh, Pittsburgh, PA, USA; Fondazione RiMED, Palermo, Italy; Department of Bioengineering, University of Pittsburgh, Pittsburgh, PA, USA; Department of Surgery, University of Pittsburgh, Pittsburgh, PA, USA

**Keywords:** hydrogel, myocardial infarction, porous structure, extracellular matrix

## Abstract

A variety of hydrogels have been developed for intramyocardial injection therapy after myocardial infarction. Some of these biomaterials have incorporated bioactive substances that promote local tissue regeneration and integration, while others have emphasized the mechanical role of the injectate in providing functional benefit. In this study, we incorporated two bioactive features, porosity, and extracellular matrix derived hydrogel (ECM), into a mechanically optimized, thermoresponsive, degradable hydrogel (poly(N-isopropylacrylamide-co-N-vinylpyrrolidone-co-MAPLA) copolymer) and evaluated whether injection of this biomaterial could abrogate the remodeling process in a rat ischemic cardiomyopathy model. After myocardial infarction by coronary artery ligation, rats were randomly divided into four groups: group NP (non-porous hydrogel) without either ECM or porosity, group PM (porous hydrogel) from the same synthetic copolymer with mannitol beads as porogens, and group PME with porosity and ECM digest added to the synthetic copolymer. A group with PBS injection alone served as a control. Intramyocardial injections were made 3 days after myocardial infarction. Serial echocardiography was assessed over time, and histological assessments were performed eight weeks after infarction. Results demonstrated improved echocardiographic function and neovascularization in the PME group compared to the other hydrogels and PBS injection. The PME group also demonstrated significant improvement in LV geometry and macrophage polarization (towards M2) compared to the PBS group, whereas these differences were not observed in the NP or PM groups versus the control. These results demonstrate that further functional improvement may be achieved in hydrogel injection therapy for ischemic cardiomyopathy by incorporating porosity and ECM digest, representing a combination of mechanical and biological effects.

## 1. Introduction

Ischemic heart disease (IHD), a major source of morbidity and mortality worldwide^1^, results when infarcted myocardium from the obstruction of coronary arteries leads to ischemic cardiomyopathy (ICM). This condition can commonly progress to end-stage heart failure (HF). The mortality of HF remains high (20.2 % and 52.6 % at 1 and 5 years after diagnosis, respectively) even with current optimal therapies.^2^ The progressive clinical deterioration after myocardial infarction (MI) is driven by a remodeling process in the left ventricle (LV). This process positively compensates cardiac function in the acute phase, but chronically contributes to irreversible LV dysfunction and structural deformation.^3^ ICM is characterized by progressive wall thinning and dilation of the infarcted ventricular segments coupled with an elevation in wall stress. The wall thinning and dilatation during the adverse remodeling increases the wall tension, which triggers further LV deformation, spiraling down a pathway towards cardiac decompensation and death.^4^

Intramyocardial injection therapy post-MI has been widely investigated during the past two decades. While some injectates have focused principally on incorporating biologically active components, others have aimed to reduce LV wall stress by injecting mechanically stiffer agents to reduce the mechanical load in the infarct or border zone region, thereby interrupting or slowing the adverse remodeling process. We and others have designed a variety of hydrogels for this purpose and have shown their therapeutic effects in animal models.^5–10^ One of these injectates that we have pursued is a group of poly(N-isopropylacrylamide)(PNIPAAm)-based hydrogels that incorporate variously optimized features, including temperature-dependent stiffening upon injection, tunable mechanical properties pre- and post-injection, and controllable degradation in situ.^11–13^ Optimized versions of this hydrogel have demonstrated functional and geometrical improvements on LV remodeling in a porcine model of ICM.^7^

To further interrupt the adverse remodeling response post-MI, and to promote retention of cardiac function and geometry, we have focused on incorporating bioactive functionality into these mechanically promising biomaterials.^14^ To this end a thermoresponsive PNIPAAm composite hydrogel was developed where D-mannitol particles were used as a porogen, with the expectation that the presence of porosity would stimulate further tissue ingrowth and elaboration in the infarcted ventricular wall. The polymer design was extended by the addition of both soluble porogen and extracellular matrix (ECM) digest hydrogel derived from porcine urinary bladder matrix.^6^ Using these three hydrogel designs (non-porous, porous, and porous + ECM), we previously demonstrated that in a rat hindlimb intramuscular injection model, porous hydrogels induced more rapid cellular infiltration and further addition of ECM content resulted in greater M2 macrophage polarization in the infiltrate compared to control groups not incorporating ECM. Based on this experience, our objective in the current report was to investigate the hypothesis that hydrogels with bioactive ECM and porous structure would decrease the infarction size, preserve cardiac wall thickness and improve cardiac function in a rat MI model by altering the adverse remodeling process in the infarcted left ventricular wall.

## 2. Materials and Methods

### 2.1 Materials

All chemicals were purchased from Sigma-Aldrich (St. Louis, MO, USA), unless otherwise specified. Mannitol particles (Sigma-Aldrich) were prepared by size separation between 170 and 230 meshes (63-88 μm) before use. Methacrylate polylactide (MAPLA) was synthesized as described previously.^12^ Briefly, NaOCH3/methanol was added to a lactide/dichloromethane solution to synthesize polylactide (HO-PLA-OCH3), and then MAPLA was synthesized by dropping methacryloyl chloride into an HOPLA-OCH3/dichloromethane solution with triethylamine.

Poly(N-isopropylacrylamide-co-N-vinylpyrrolidone-co-MAPLA) copolymer (poly(NIPAAm-co-VP-co-MAPLA) was synthesized from NIPAAm, VP and MAPLA by free radical polymerization as previously described,^11^ using NIPAAm, VP and MAPLA at 80/10/10 feed ratios. Monomers (0.066 mol) were dissolved in 1,4-dioxane (180 mL) with 0.23 g BPO. Polymerization proceeded at 70 C for 24 h under an argon atmosphere. The copolymer was precipitated in hexane with purification by precipitation from THF into diethyl ether and vacuum-drying, to provide ∼80% yields.

Using a published technique, ECM digest hydrogel was prepared from urinary bladder matrix (UBM) digest as previously described.^15^ Briefly, fresh porcine bladders (Thoma Meat Market, Pittsburgh, PA) were cleaned and excess connective tissue excluded, followed by mechanical removal of the tunica serosa, tunica muscularis externa, the tunica submucosa, and the tunica muscularis mucosa. The luminal surface was rinsed with 1.0 N saline to dissociate urothelial cells of the tunica, leaving basement membrane and subadjacent lamina propria, or UBM. Sheets of UBM were placed in a 0.1% (v/v) peracetic acid solution, 4 % (v/v) ethanol, and 95.9 % (v/v) sterile water for 2 h followed by 2 x 15 min PBS rinses and 2 x15 min washes with sterile water. The sheets were lyophilized and milled into powder and filtered through a 60 mesh screen. Powdered UBM was solubilized at 10 mg/mL in 0.1 mg/mL pepsin in 0.01 N HCl at a constant stir rate for 48 h and then neutralized to pH 7.4 with NaOH and diluted in phosphate buffered saline (PBS).^15^

Three different hydrogel types were prepared for later injection studies: 1) nonporous hydrogel (NP) containing 10 w/v% copolymer in PBS, 2) a porous hydrogel (PM) composed of 10w/v% copolymer with 30 w/v% sized mannitol particles, and 3) a porous hydrogel with UBM digest (PME) with 20w/v% copolymer mixed with 10 mg/mL UBM digest mixture (1:1, v/v), followed by addition of 30 w/v% sized mannitol particles. All samples were prepared and stored at 4°C prior to use.

### 2.2 Animal model

The 10- to 12-week-old adult female syngeneic Lewis rats (ENVIGO, Indianapolis, IN, USA) weighing 180-220 g were used for this study. The research protocol followed the National Institutes of Health guidelines for animal care and was approved by the Institutional Animal Care and Use Committee of the University of Pittsburgh (#20097929). Rats were randomly divided into four groups reflecting the injectate used post-MI: group NP without either ECM or pores (n = 7), group PM with pores (n = 8), group PME with pores and ECM (n = 8), and a control group with PBS injection (n = 8) (**Fig. 1**).

**Fig. 1.**
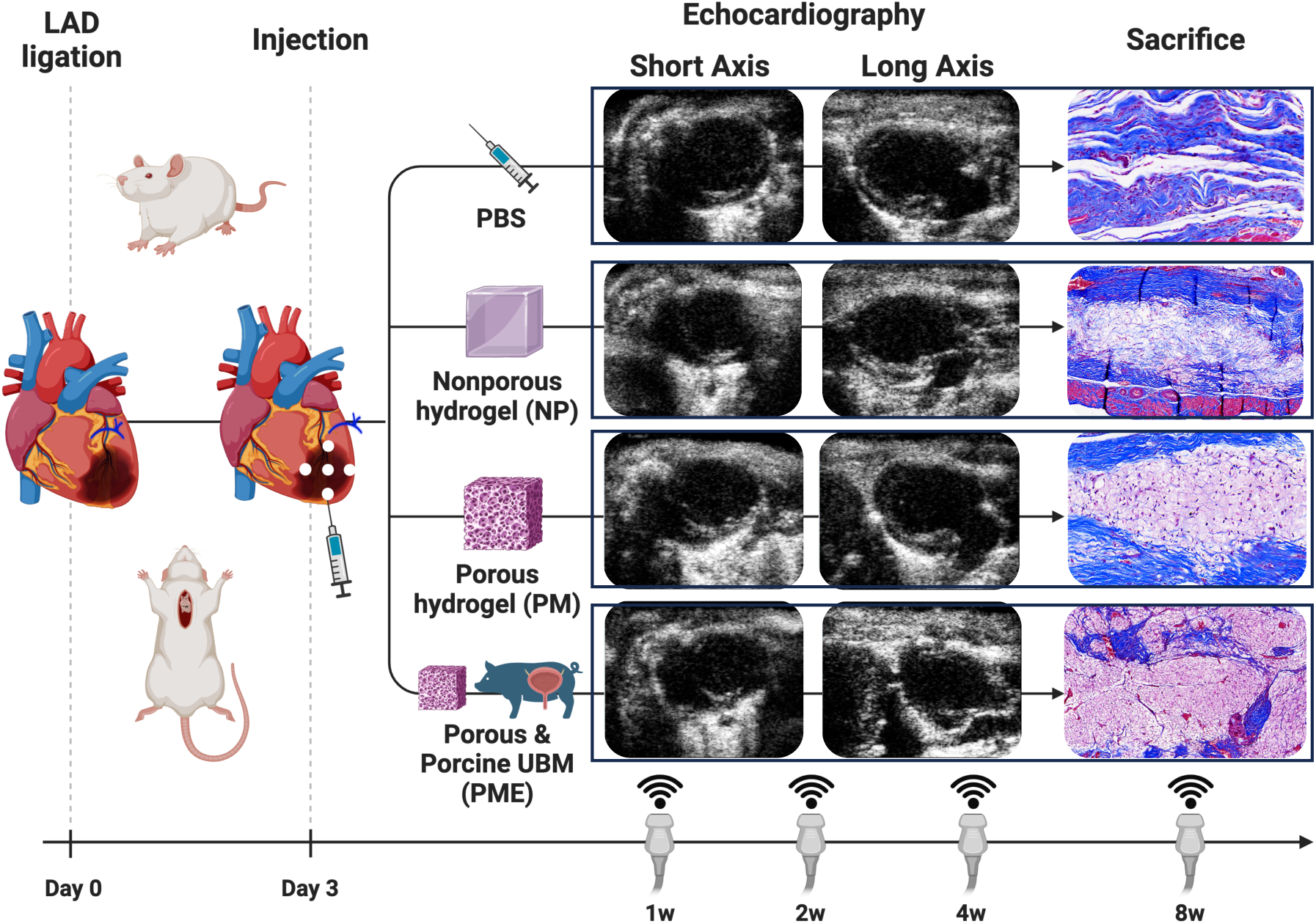
Timeline of study. All echocardiographic images in this figure were taken during the diastolic phase at 8 weeks before explant. The Masson’s trichrome staining images show the injection sites.

### 2.3 Procedures for MI model creation and hydrogel injection

Left myocardial infarction was created by ligation of the proximal left anterior descending artery (LAD) as previously described.^16^ Briefly, the rat was anesthetized with 2.5 % isoflurane induction and 1.25 to 1.5 % maintenance with 100 % oxygen followed by intubation and respiratory support with a rodent volume-controlled mechanical ventilator (683 Ventilator, Harvard Apparatus, Holliston, MA, USA) at a tidal volume of 2 mL and 60-70 breaths/min. The rat was placed in the supine position on a warming blanket (37 ℃), and the chest was shaved and prepared with povidone-iodine solution. Before skin incision, 10 mg/kg lidocaine hydrochloride as an analgesic and an antiarrhythmic agent and cefazolin as an antibiotic were administered intramuscularly. All procedures were performed in a sterile environment. The heart was exposed through the 4th left thoracotomy, and the proximal LAD was directly ligated with 5-0 polypropylene. The incision was closed with 4-0 continuous sutures. The animals were allowed to recover from anesthesia and returned to their cages. The hydrogel injection was performed 3 days after infarction creation as previously described.^7^ The rat was anesthetized and screened by echocardiography for infarct size in terms of the percentage of scar area (akinetic or dyskinetic regions) to LV free wall area. Only rats with infarction >25 % of the LV free wall were chosen for further procedures. At the time of injection, the infarcted rat heart was exposed through the same left thoracotomy, and hydrogel or PBS was injected into infarct border zones and the center of the infarct (5 injections, 20 µL per region, 100 µL in total). The volume of injected hydrogel was determined based on the surgeon’s experience in injecting biomaterials into this rat model to maximize the volume injected in general.

### 2.4 Wall thickness and scar size in LV anterior wall

The cross sections with Masson’s trichrome staining were digitally photographed. The wall thickness was expressed as follows: Area of region of interest / [(epicardial circumference + endocardial circumference)/2]. We measured wall thicknesses in the injection site inside the myocardial infarcted area as well as the whole myocardial infarcted area. The scar area was defined as a blue-stained area (with Masson’s trichrome staining), and the scar size (%) was calculated by dividing the scar area by the total left ventricular area. Each parameter was measured using ImageJ (v.1.54b, National Institutes of Health, Bethesda, MD, USA).

### 2.5 Echocardiography

Cardiac functional parameters were evaluated with echocardiography before surgery, 3 days after MI, and 1, 2, 4, and 8 weeks after hydrogel injection. Rats were anesthetized with 1.5-2.0 % isoflurane inhalation with 100 % oxygen without mechanical ventilation. Transthoracic echocardiography was performed using the Acuson Sequoia C256 system with 13-MHz linear ultrasonic transducer (15L8; Acuson Corporation, Mountain View, CA, USA) in a phased array format. Left ventricular (LV) parameters recorded were the end-diastolic area (EDA), end-systolic area (ESA), end-diastolic dimension (LVDd) and end-systolic dimension (LVDs) as obtained from the short axis view at the papillary muscle level. The LV fractional area change (%FAC) and fractional shortening (%FS) were calculated as %FAC = (EDA-ESA)/EDA x 100 % and %FS = (LVDd-LVDs)/LVDd x 100 %, respectively. Ventricular volume (V) was estimated using the formula of Teichholz to calculate LV end-diastolic volume (LVEDV) and end-systolic volume (LVESV) as follows: V = 7.0/(2.4+D) x D³, where D is the LV diameter measured by M-mode echocardiography. LV ejection fraction (LVEF) was calculated as LVEF = (LVEDV–LVESV)/LVEDV x 100 %.

### 2.6 Histology and immunohistochemistry

Harvested heart tissues were fixed with 10 % buffered formalin and embedded in paraffin. Five um-thick serial paraffin-embedded sections were deparaffinized in xylene, dehydrated in graded ethanol mixtures, and stained with hematoxylin and eosin, and Masson’s trichrome. Paraffin-embedded sections were blocked with staining buffer for 1 h (10 % goat serum with 1 % BSA in PBS). For macrophage analysis, the primary antibodies were rabbit anti-CD68 antibody (ab125212, 1:100, Abcam, Cambridge, MA), mouse anti-CD86 antibody (ab220188, 1:100, Abcam), and mouse anti-CD206 antibody (sc-58986, 1:100, Santa Cruz, Dallas, TX). The secondary antibodies were a goat anti-rabbit IgG Alexa Fluor 568 (1:200) for pan-macrophages and a goat anti-mouse IgG Alexa Fluor 488 (1:200) for M1 and M2 macrophages. For vessel analysis, the primary antibodies were a mouse anti-α-SMA antibody (ab7817, 1:200, Abcam) and a rabbit anti-Von Willebrand factor (vWF) antibody (ab6994, 1:200, Abcam). The secondary antibodies were a goat anti-mouse IgG Alexa Fluor 488 (ab150117, 1:500, Abcam) for α-SMA, and a goat anti-rabbit IgG Alexa Fluor 568 (ab175695, 1:500, Abcam) for vWF. Nuclei were stained with 4’,6-diamidino-2-phenylindole dihydrochloride (DAPI, ab104139, Abcam). Multispectral epifluorescent images were acquired using a Nikon Eclipse 6600 microscope (Nikon corporation, Tokyo, Japan), with spectral unmixing to remove autofluorescence using Nuance 3.0.2 software (Caliper Life Science Inc., Hopkinton, MA). To quantify macrophage infiltration, M1 macrophages were defined as CD68/CD86 double-positive cells and M2 macrophages were defined as CD68/CD206 double-positive cells, identified using digital image analysis software (CellProfiler v.4.2.1, Broad Institute, inc., Cambridge, MA). To quantify vessel distribution in the injection sites, each vessel was identified as a tubular structure with a visible lumen of > 10 µm in diameter, positively stained for α-SMA and vWF in the infarction sites. The total percent area occupied by blood vessels (the vessel area %) and the number of vessels were quantified in the region of interest (microscope field). The vessel measurements were performed using ImageJ.

### 2.7 Statistical analyses

One-way repeated measures analysis of variance (ANOVA) followed by Tukey’s test was applied for all multi-group comparisons in the study. All statistical analyses were conducted using GraphPad Prism for Mac (Version 8, San Diego, CA, USA). Data are expressed as mean ± standard error of the mean (SEM). P-values < 0.05 were considered significantly different.

## 3. Results

### 3.1 LV wall histology after infarction

The Masson’s trichrome stained images show the infarcted and scarred LV wall at 8 wk post-MI (**Fig. 2A**). The magnified images of Fig. 2A show detail of the injection sites in MI areas (**Fig. 2B**). The wall thickness in the whole MI area (**Fig. 2C**), and injection site inside the MI area (**Fig. 2D**) were larger in the NP, PM, and PME groups than the PBS group. There were also significant differences in the scar size as a percentage of the total left ventricular area between the therapeutic groups and the control (**Fig. 2E**).

**Fig. 2.**
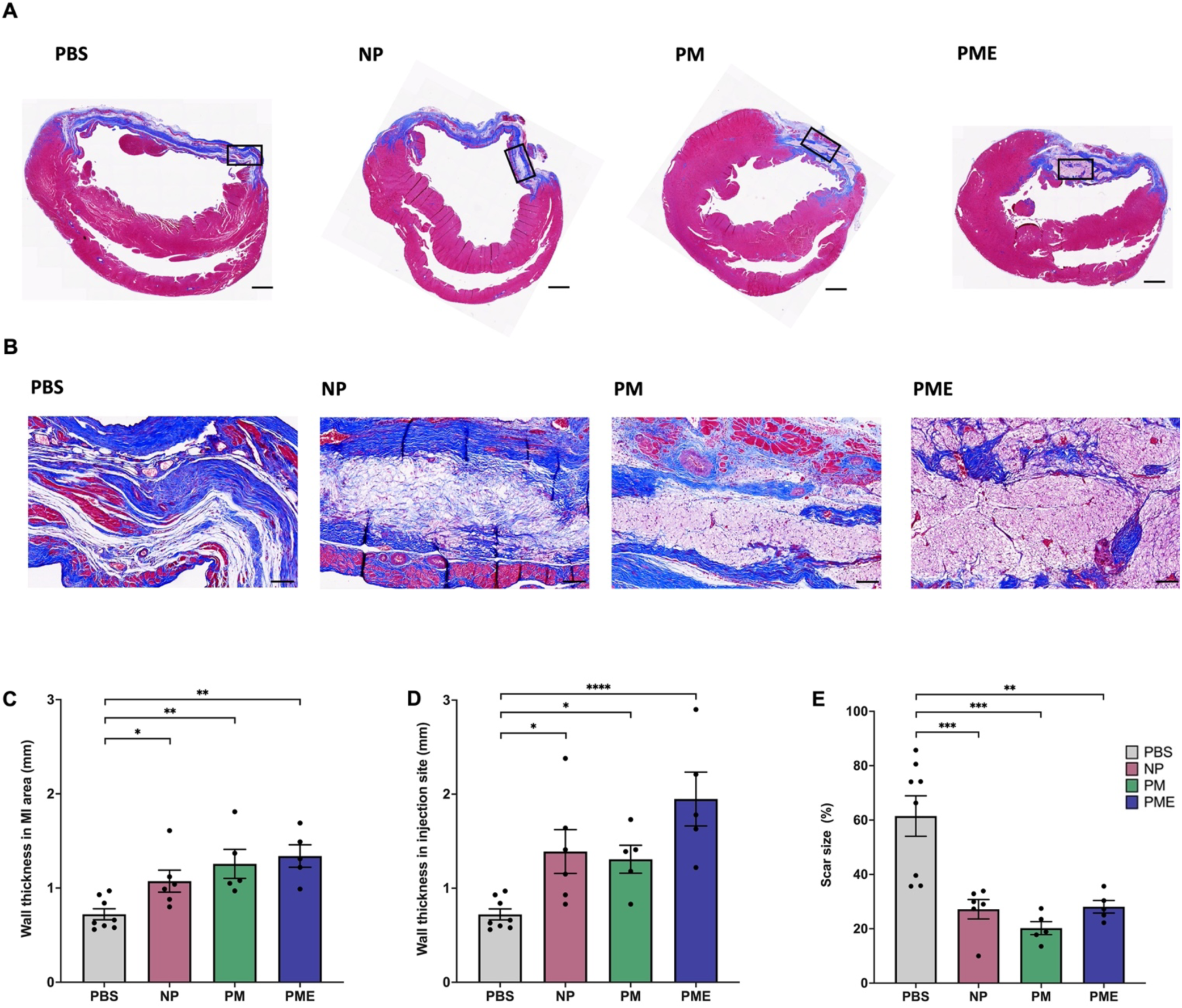
Left ventricular wall thickness and scar size. The representative images from each group of the left ventricular wall with Masson’s trichrome staining. Scale bars = 1 mm. (B) The zoom images of box regions in (A), focusing on and around injection sites. Scale bars = 100 μm. The LV wall thickness (C) in the myocardial infarction area and (D) in injection sites. (E) The scar size in the myocardial infarction area. The PBS (n = 8), NP(n = 6), PM (n = 5), and PME (n = 5). *p < 0.05, **p < 0.01, ***p < 0.001, ****p <0.0001.

### 3.2 Echocardiography

The results of echocardiography before surgery, 3 days after MI, and 1, 2, 4, and 8 weeks after hydrogel injection are shown in **Fig. 3**. There were no differences in all parameters of cardiac function between the groups before surgery. At 8 weeks, the geometrical analyses showed that PME was significantly smaller in LVDd, LVDs, EDA, ESA, EDV, and ESV than the PBS group (**Fig. 3A-F**). There were no significant differences in geometrical parameters between PME and the other hydrogel groups (PM or NP). On the other hand, in the functional assessments at 8 weeks, PME showed greater function in %FS and LVEF compared to NP and PM, and greater %FS, %FAC, and LVEF compared to PBS (**Fig. 3G-I**).

**Fig. 3.**
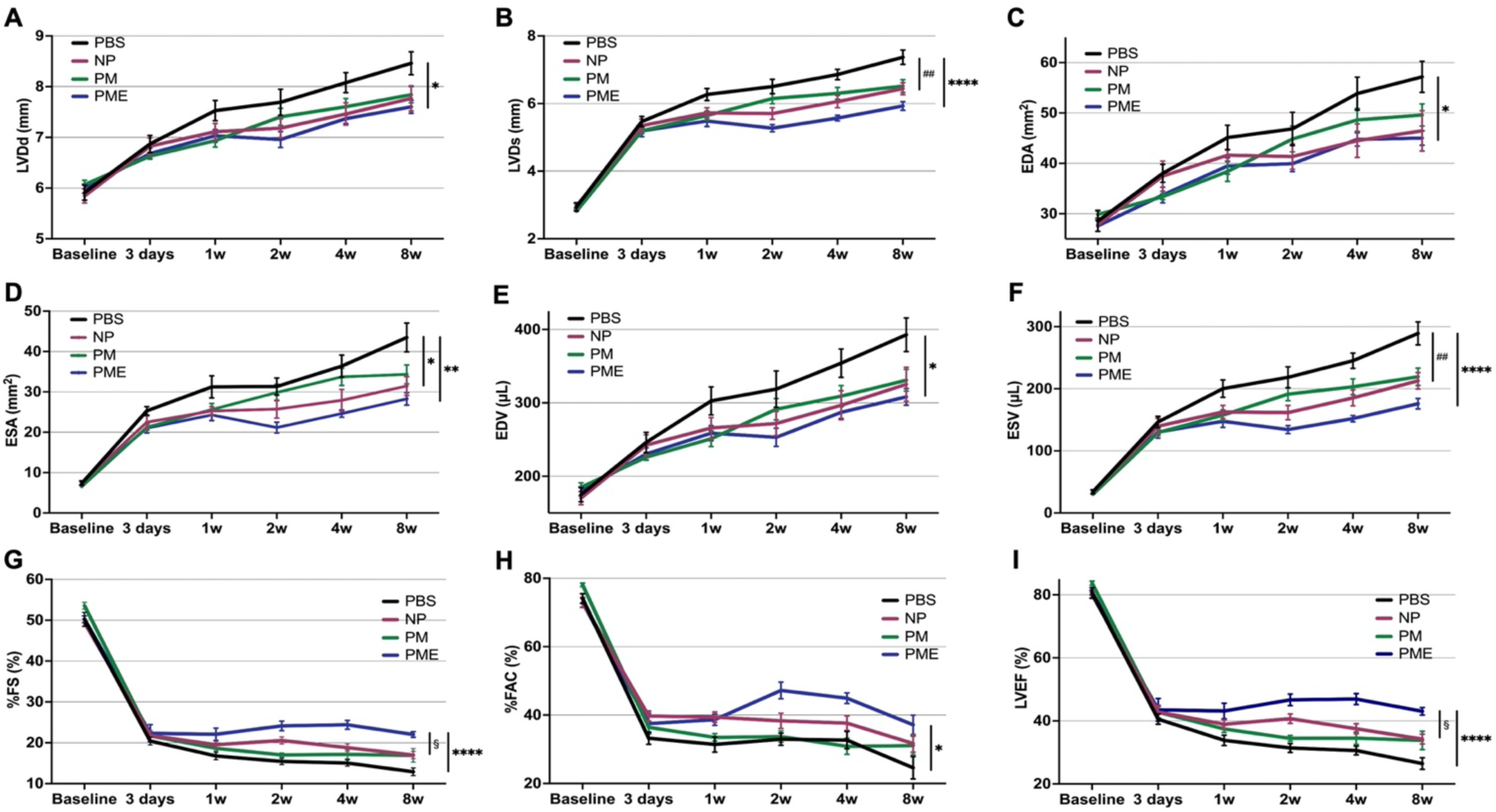
Echocardiography. (A) Left ventricular end-diastolic diameter (LVDd), (B) Left ventricular end-systolic diameter (LVDs), (C) End-diastolic area (EDA), (D) End-systolic area (ESA), (E) End-diastolic volume (EDV), (F) End-systolic volume (ESV), (G) Fractional shortening (%FS), (H) Fractional area change (%FAC), (I) Ejection fraction (EF). All data are means ± SEM and were assessed by one-way ANOVA followed by Tukey’s multiple comparisons test. *p < 0.05, **p < 0.01, and ****p < 0.0001 between the PME and PBS groups. ##p < 0.01 between the PM, NP, and PBS groups. §p < 0.05 between the PME and the other hydrogel groups (the PM or NP).

### 3.3 Macrophage infiltration and polarization

As shown in **Fig. 4A-D**, staining for CD68, CD86, and CD206 showed the distribution of CD68 single-positive cells (representing all macrophages), CD86/CD68 double-positive cells (consistent with M1 macrophages), and CD206/CD68 double-positive cells (consistent M2 macrophages). The percent ratio of CD86/CD68 double-positive cells (M1 macrophages) to CD68 single-positive cells (all macrophages) in the ischemic areas was lower in the PM and PME groups than in the control group (**Fig. 4E**). Whereas the percent ratio of CD206/CD68 double-positive cells (M2 macrophages) to CD68 single-positive cells (all macrophages) in the injection areas was higher in the NP, PM, and PME groups than in the control group (**Fig. 4F**). The ratio of CD206/CD68 double-positive cells (M2 macrophages) to CD86/CD68 double-positive cells (M1 macrophages) in the injection area was higher in the PME group than in the control group (**Fig. 4G**).

**Fig. 4.**
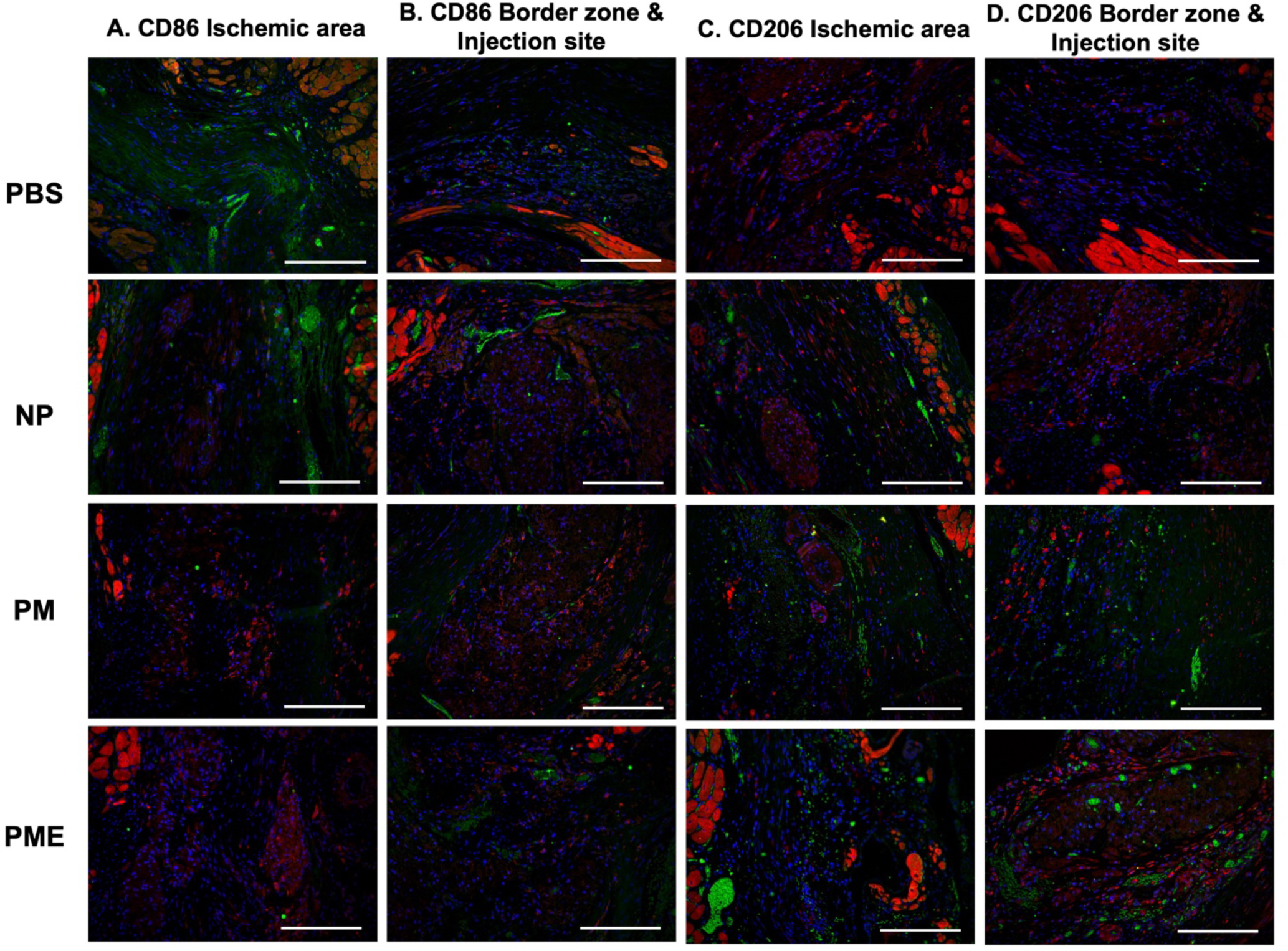
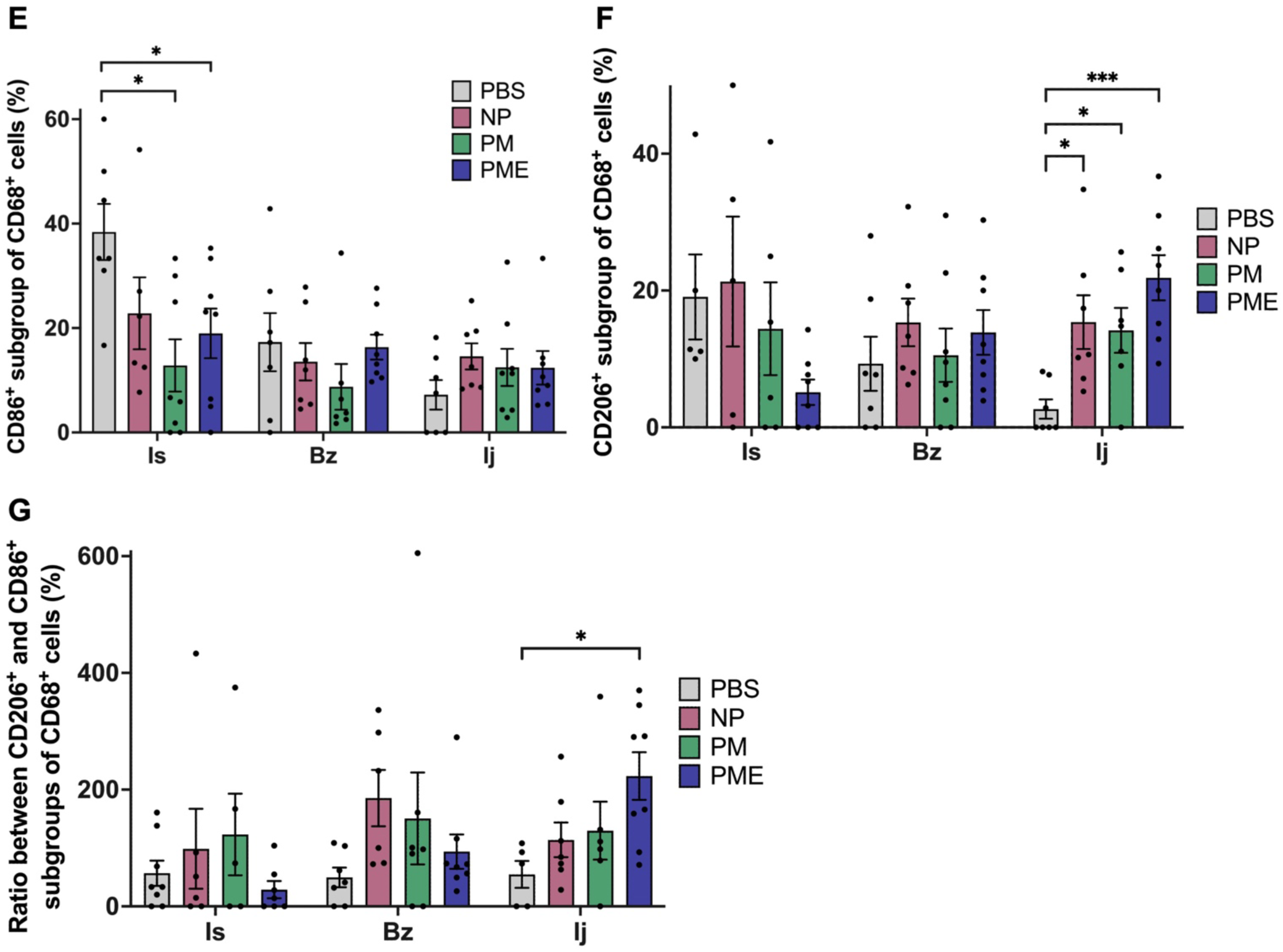
Macrophage infiltration and polarization. The representative images of immunofluorescence staining for (A, B) CD68 (red) and CD86 (green), and (C, D) CD68 (red) and CD206 (green) in the ischemic areas, and injection sites & injection border zones. Red-stained areas that appear to be myocardium structures were excluded as autofluorescence. All scale bars = 100 μm. (E) The percent ratio of M1 macrophages to all macrophages in the ischemic areas. (F) The percent ratio of M2 macrophages to all macrophages in the injection areas. (G) The ratio of M2 macrophages to M1 macrophages in the injection area. Data are means ± SEM. *p < 0.05 and ***p < 0.001 assessed by one-way ANOVA followed by Tukey’s multiple comparisons test. Is: Ischemic area, Bz: injection border zone, Ij: injection site.

### 3.4 Neovascularization

Immunofluorescence staining for α-SMA and vWF was performed to illustrate the vessel distribution in the ventricular wall at 8 weeks (**Fig. 5A, B**). As shown in **Fig. 5C**, the vessel area in injection sites was higher in the hydrogel groups than in the PBS group. Furthermore, the PME group showed a higher vessel area than the NP group in the ischemia area and border zone but not in the injection site. The number of vessels per field was significantly higher in the hydrogel groups than in the control group in the ischemic area and injection site. The PME group showed more vessels than the NP group in all three areas and the PM group in the ischemic area (**Fig. 5D**).

**Fig. 5.**
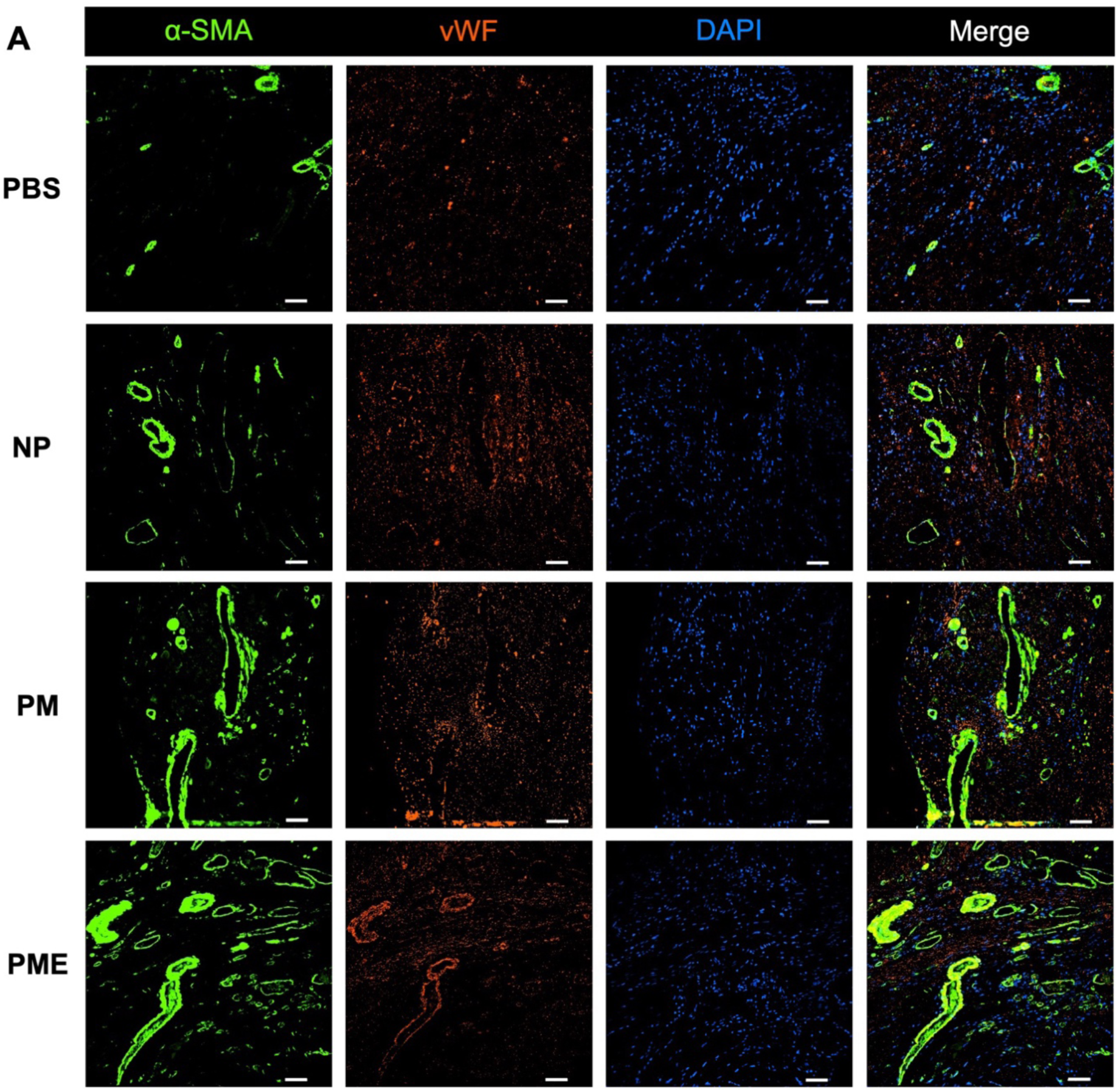
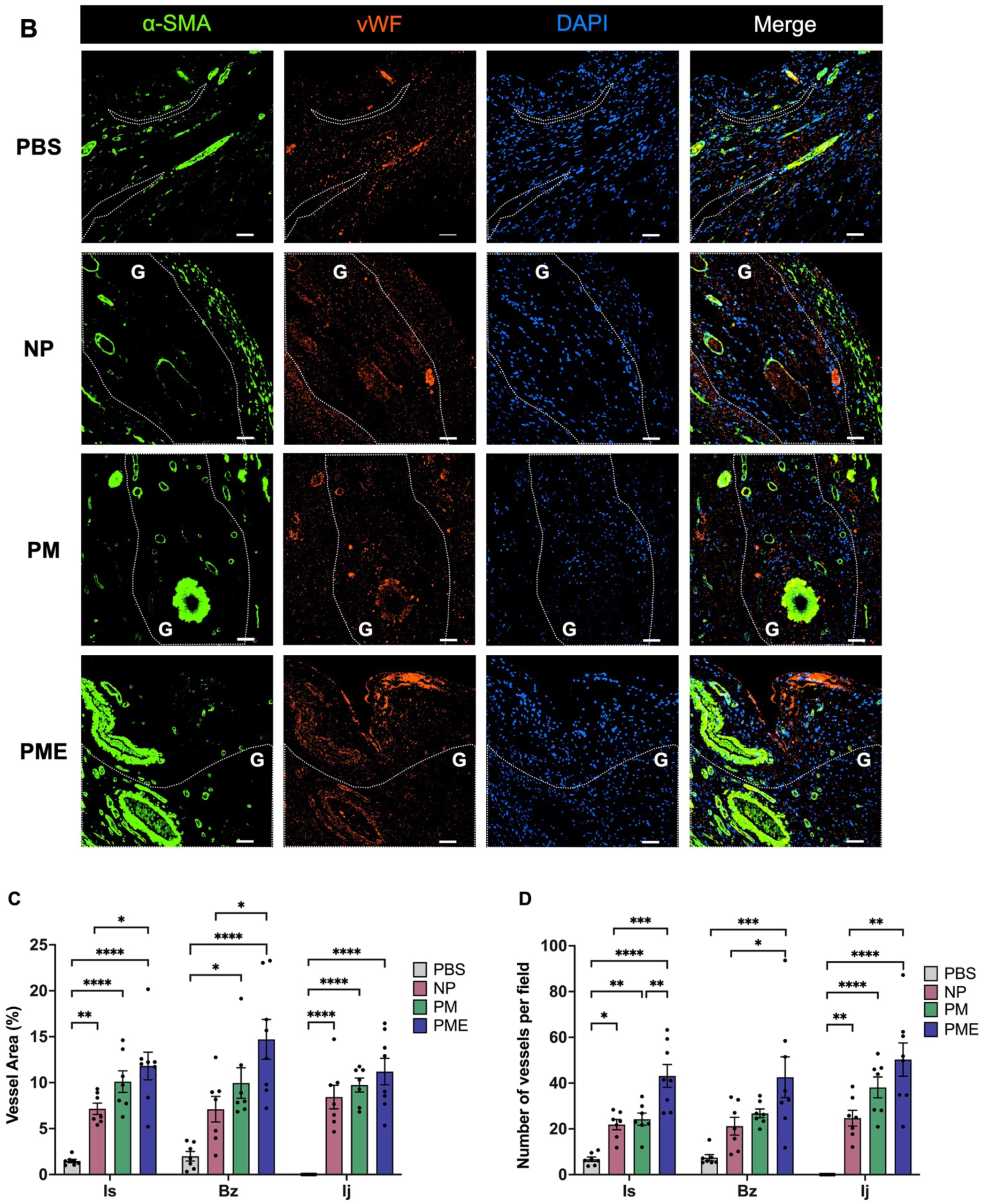
Vessel analysis. The representative images of immunofluorescence staining for α-SMA (green) and vWF (red) in (A) ischemic areas and (B) border zones and injection sites. The areas encircled with dashed white lines correspond to the injection sites. All scale bars = 50 μm. G: hydrogel. Measurements of (C) vessel density per ROI region and (D) number of vessels per field. Data are means ± SEM. *p < 0.05, **p < 0.01, ***p < 0.001, and ****p < 0.0001 assessed by one-way ANOVA followed by Tukey’s multiple comparisons test. Is: Ischemic area, Bz: injection border zone, Ij: injection site.

## 4. Discussion

This study examined the response to three novel thermoresponsive hydrogels in a rat ICM model with an open chest injection approach. A similar NIPAAM based thermoresponsive hydrogel with MAPLA segments has already demonstrated its positive effects in preserving cardiac function and structure in a previous report using a porcine MI model.^7^ We designed this hydrogel to have comparable features, in terms of injectability followed by rapid stiffening in situ and degradation.^6^ Based on this optimized hydrogel, in this report the functionality of the hydrogel was expanded by adding porosity and an ECM component, with hypothesized additional biological effects. Supporting this hypothesis, more pronounced effects in preserving LV function were seen in the PME group compared to the other hydrogels and the PBS group, specifically in %FS and LVEF. The PME group preserved all LV geometrical parameters relative to the PBS group, whereas differences were only observed in LVDs and ESV for the NP or PM groups versus the control. Regarding the ratio between M2 and M1 macrophages, the PME group had higher levels of this ratio than the PBS group in the injection site, whereas no differences were observed for the NP or PM groups versus the control. Moreover, the PME group consistently had greater levels of vascularization and vessel area relative to the other two hydrogel groups and PBS controls.

Despite various efforts and treatment strategies, there is still no definitive treatment in preventing the onset or progression of ICM resulting from adverse LV remodeling. To remedy end-stage HF, various surgical strategies have been attempted, such as dynamic cardiomyoplasty^17, 18^ and ventricular restraint therapies^19^. However, most have been withdrawn from clinical practice or are only applied in limited patient populations today. Alternatively, coronary revascularization and optimal medical therapy have expanded in recent years.^20^ The limitation of revascularization therapy is that the success largely depends on how many viable cardiomyocytes can be salvaged, because lost myocardium cannot be regained with this therapy. Heart transplantation or ventricular assist devices are applied as the last options for patients with end-stage HF. Still, requests for organ transplantation continue to substantially outpace the number of donor organs^21^, and highly invasive mechanical support carries acute and chronic risks from infection, thromboembolism and bleeding as well as lifestyle compromises.^22^

There is a long interval in treatment between the period when revascularization could provide beneficial effects and the period when a patient has reached end-stage HF, representing both a need and opportunity for more advanced therapies. Treatment that preserves or restores cardiomyocytes in the face of adverse remodeling during this period is desirable to fill this gap. Many studies have evaluated stem cell therapy approaches in recent years, aiming to regenerate and restore functioning cardiomyocytes. However, the outcomes thus far have been modest considering the original aims due largely to the low cell retention rate in impaired tissue. Today, this therapy is believed to contribute primarily through paracrine effects.^23–25^

### 4.1 Stiffness

Elastic modulus is a tunable parameter, and the stiffness of injected hydrogels is believed to be an essential factor in modulating LV remodeling. The stress reduction in the infarct border zone region may contribute to minimizing stress-induced apoptosis and infarct border zone expansion, suppressing further adverse remodeling.^26^ The modulus of human myocardium is 10–20 kPa during diastole, and 200–500 kPa during systole, which is much stiffer than alginate hydrogel (3–5 kPa) and ECM-derived hydrogel (2–15 Pa) that have been used in the clinical trials.^27–29^ The relatively low modulus might be one of the reasons why these clinical trials have failed to demonstrate adequate therapeutic effects despite encouraging pre-clinical results in large animal models.^30–32^ Higher stiffness hydrogels are recommended in myofiber stress reduction according to some finite element (FE) model simulations.^33, 34^ As a landmark study in this field, Wall et al. showed that stiffer materials with a higher elastic modulus efficiently improved local wall stresses and LV metrics with a FE model.^33^ In later studies with animal models, the efficacies of higher stiffness hydrogels have also been demonstrated in terms of myofiber stress reduction, smaller infarct area, and ejection fraction.^35–37^ More specifically, another FE model incorporating experimental data showed that diastolic myofiber stress was more effectively reduced as the stiffness reached higher values in a range of 5 kPa to 100 kPa, but this beneficial effect tapered after 50 kPa.^38^ One of the drawbacks of ECM hydrogel is its relatively low modulus in terms of mechanical support for the infarcted LV.^29, 32^ The compression modulus of hydrogel NP in our present study was 272 ± 38 kPa, and the compression modulus of the PME was 87 ± 6 kPa despite the presence of ECM and the porous structures inside the hydrogel, still falling into the range of the simulated requirements above.^6^

### 4.2 Porosity

It remains a significant challenge in synthetic hydrogel development to effectively integrate bioactive functions promoting tissue regeneration. In recent years, the concept that controlled porosity in hydrogels could better promote tissue integration and vascularization has been a research focus.^39, 40^ Parameters such as pore size and distribution, porosity and interconnectivity are important in facilitating cell behaviors, such as proliferation,^41^ differentiation^42^, and adhesion^43^. Moreover, the presence of pores in a scaffold generally reduces stiffness as pore size and porosity increase, which would be counter to the stiffness benefit from the theory of Wall et al.^33^ The optimal pore size from the perspective of cellular behavioral regulation depends on which cells or tissues are targeted.^42–44^ It has been reported that different pore sizes in implants affects macrophage polarization,^40^ with macrophages showing a transition from M1 towards M2 phenotype around implant pores. However, the influences of porosity and pore profile on remodeling after MI have not yet been well studied. In the present report a highly soluble mannitol bead was used as a porogen. While the porogen was sieved to obtain beads sized between 63-88 μm, the formed pore sizes of PM and PME were 237 ± 80 μm and 295 ± 87 μm, respectively.^6^ The parameters for mannitol bead size and added amount were based on an earlier report in skeletal muscle and the demonstration that the selected values in that study provided extensive cellular infiltration and an M2 versus M1 effect. Although the stiffness was reduced with the introduction of porosity, PM and PME injectates still preserved enough compressive modulus to support the LV wall mechanically.^6^

### 4.3 ECM

Other efforts to acquire bioactive functions include incorporating bioactive substances directly into the hydrogels. ECM is naturally bioactive and plays an important role in the inflammation and remodeling process after MI via cell signaling processes.^45^ ECM-derived hydrogels with inherent bioactivity have been hypothesized to modulate the inflammatory responses and positively impact adverse remodeling. Decellularized tissue, processed into an injectable hydrogel form, is comprised of remnants of tissue ECM, a complex network of proteins, proteoglycans, and glycosaminoglycans. ECM hydrogel provides not only adhesive sites for infiltrating cells but also bioactive cues to endogenous cells promoting cellular infiltration and differentiation, which is expected to play an essential role in the inflammation and remodeling process after MI.^45^ An emulsion derived from small intestine ECM was associated with functional improvements following post-MI injection with recruitment of macrophages.^46^ M2 macrophage polarization, the transition from the pro-inflammatory M1 phenotype to the constructive and modulatory M2 phenotype, has also been recognized in recent years.^47^ Some ECM-derived biomaterials have shown positive outcomes in MI therapy. VentriGel^TM^, derived from decellularized porcine myocardial tissue, demonstrated increased cardiac muscle, reduced fibrosis, and significant improvements in cardiac function in a porcine MI model.^48^ In the subsequent phase I clinical trial, this hydrogel also showed improvements in some clinical parameters, including 6-min walk test distance and New York Heart Association functional class.^32^

### 4.4 Limitations

This study has several limitations. To inject the hydrogels, a conventional open chest approach was used. This was done since the rat model was too small to apply alternative less-invasive approaches. However, such minimally invasive approaches are a theoretical benefit of injection therapy and would need to be demonstrated in a large animal model. Also, the volume and distribution of hydrogel injection is difficult to control and vary with the small animal heart. Hydrogel injection might be optimized based on the specific location of the infarct. Specifically, recent finite element mechanical models employ clinical measurements from MRI and catheterization results to create sophisticated simulations.^49, 50^ These in silico models with location data regarding the infarcted segments obtained from MRI could provide guidance for an optimal injection pattern for a given infarcted ventricle. This might lead to further benefits as injection therapy is translated to the clinic.

## 5. Conclusion

In this study, we aimed to show the potential therapeutic effects associated with a thermoresponsive hydrogel incorporating porosity and ECM components in a rat ICM model. Our results indicate that adding porosity together with ECM had an additional positive effect on cardiac function over hydrogel alone. The PME group also significantly improved LV geometry compared to the PBS group in all parameters, whereas the NP and PM groups were improved in only a few parameters. In terms of inflammatory response, the ratio between M2 and M1 macrophages was greater in the PME group than the PBS group in the injection site, whereas no differences were observed for the other hydrogels versus the control. There was also consistently greater levels of vascularization and vessel area with porosity and ECM content versus the other two hydrogel groups and PBS controls. The results support the advancement of this hydrogel design toward clinically relevant large animal studies, which would allow the utilization of minimally invasive approaches and potentially infarct mechanical modeling to optimize injection strategies.

## Ethics approval statement

The Institutional Animal Care and Use Committee of the University of Pittsburgh (#20097929) reviewed and approved the animal study.

## Author Contributions

Y.H. and T.F.: conceptualization, data curation, formal analysis, investigation, methodology, writing-original draft, writing-review & editing. S.K., S.B. and A.D.: resources; writing-review & editing. W.W.: conceptualization, funding acquisition; supervision; writing-review & editing. Y.H. and T.F. contributed equally to this work. All authors contributed to the article and approved the submitted version.

## REFERENCES

1. Martin SS, Aday AW, Almarzooq ZI, Anderson CAM, Arora P, Avery CL, Baker-Smith CM, Barone Gibbs B, Beaton AZ, Boehme AK, Commodore-Mensah Y, Currie ME, Elkind MSV, Evenson KR, Generoso G, Heard DG, Hiremath S, Johansen MC, Kalani R, Kazi DS, Ko D, Liu J, Magnani JW, Michos ED, Mussolino ME, Navaneethan SD, Parikh NI, Perman SM, Poudel R, Rezk-Hanna M, Roth GA, Shah NS, St-Onge MP, Thacker EL, Tsao CW, Urbut SM, Van Spall HGC, Voeks JH, Wang NY, Wong ND, Wong SS, Yaffe K, Palaniappan LP. 2024 Heart Disease and Stroke Statistics: A Report of US and Global Data From the American Heart Association. Circulation. 2024;149(8):e347–e913. Epub 20240124. doi: 10.1161/cir.0000000000001209. PubMed PMID: 38264914.

2. Gerber Y, Weston SA, Redfield MM, Chamberlain AM, Manemann SM, Jiang R, Killian JM, Roger VL. A contemporary appraisal of the heart failure epidemic in Olmsted County, Minnesota, 2000 to 2010. JAMA Intern Med. 2015;175(6):996–1004. doi: 10.1001/jamainternmed.2015.0924. PubMed PMID: 25895156; PMCID: PMC4451405.

3. Pfeffer MA, Braunwald E. Ventricular remodeling after myocardial infarction. Experimental observations and clinical implications. Circulation. 1990;81(4):1161–72. doi: 10.1161/01.cir.81.4.1161. PubMed PMID: 2138525.

4. Mann DL, Bristow MR. Mechanisms and models in heart failure: the biomechanical model and beyond. Circulation. 2005;111(21):2837–49. doi: 10.1161/circulationaha.104.500546. PubMed PMID: 15927992.

5. Yoshizumi T, Zhu Y, Jiang H, D’Amore A, Sakaguchi H, Tchao J, Tobita K, Wagner WR. Timing effect of intramyocardial hydrogel injection for positively impacting left ventricular remodeling after myocardial infarction. Biomaterials. 2016;83:182–93. Epub 20151215. doi: 10.1016/j.biomaterials.2015.12.002. PubMed PMID: 26774561; PMCID: PMC4754148.

6. Zhu Y, Hideyoshi S, Jiang H, Matsumura Y, Dziki JL, LoPresti ST, Huleihel L, Faria GNF, Fuhrman LC, Lodono R, Badylak SF, Wagner WR. Injectable, porous, biohybrid hydrogels incorporating decellularized tissue components for soft tissue applications. Acta Biomater. 2018;73:112–26. Epub 20180410. doi: 10.1016/j.actbio.2018.04.003. PubMed PMID: 29649634; PMCID: PMC5985206.

7. Matsumura Y, Zhu Y, Jiang H, D’Amore A, Luketich SK, Charwat V, Yoshizumi T, Sato H, Yang B, Uchibori T, Healy KE, Wagner WR. Intramyocardial injection of a fully synthetic hydrogel attenuates left ventricular remodeling post myocardial infarction. Biomaterials. 2019;217:119289. Epub 20190617. doi: 10.1016/j.biomaterials.2019.119289. PubMed PMID: 31254935.

8. Chung JJ, Han J, Wang LL, Arisi MF, Zaman S, Gordon J, Li E, Kim ST, Tran Z, Chen CW, Gaffey AC, Burdick JA, Atluri P. Delayed delivery of endothelial progenitor cell-derived extracellular vesicles via shear thinning gel improves postinfarct hemodynamics. J Thorac Cardiovasc Surg. 2020;159(5):1825–35.e2. Epub 20190618. doi: 10.1016/j.jtcvs.2019.06.017. PubMed PMID: 31353103; PMCID: PMC7077034.

9. Feng J, Xing M, Qian W, Qiu J, Liu X. An injectable hydrogel combining medicine and matrix with anti-inflammatory and pro-angiogenic properties for potential treatment of myocardial infarction. Regen Biomater. 2023;10:rbad036. Epub 20230419. doi: 10.1093/rb/rbad036. PubMed PMID: 37153848; PMCID: PMC10159687.

10. Kong P, Dong J, Li W, Li Z, Gao R, Liu X, Wang J, Su Q, Wen B, Ouyang W, Wang S, Zhang F, Feng S, Zhuang D, Xie Y, Zhao G, Yi H, Feng Z, Wang W, Pan X. Extracellular Matrix/Glycopeptide Hybrid Hydrogel as an Immunomodulatory Niche for Endogenous Cardiac Repair after Myocardial Infarction. Adv Sci (Weinh). 2023;10(23):e2301244. Epub 20230615. doi: 10.1002/advs.202301244. PubMed PMID: 37318159; PMCID: PMC10427380.

11. Zhu Y, Wood NA, Fok K, Yoshizumi T, Park DW, Jiang H, Schwartzman DS, Zenati MA, Uchibori T, Wagner WR, Riviere CN. Design of a Coupled Thermoresponsive Hydrogel and Robotic System for Postinfarct Biomaterial Injection Therapy. Ann Thorac Surg. 2016;102(3):780–6. Epub 20160504. doi: 10.1016/j.athoracsur.2016.02.082. PubMed PMID: 27154150; PMCID: PMC4995147.

12. Ma Z, Nelson DM, Hong Y, Wagner WR. Thermally Responsive Injectable Hydrogel Incorporating Methacrylate-Polylactide for Hydrolytic Lability. Biomacromolecules. 2010;11(7):1873–81. doi: 10.1021/bm1004299.

13. Zhu Y, Jiang H, Ye SH, Yoshizumi T, Wagner WR. Tailoring the degradation rates of thermally responsive hydrogels designed for soft tissue injection by varying the autocatalytic potential. Biomaterials. 2015;53:484–93. Epub 20150320. doi: 10.1016/j.biomaterials.2015.02.100. PubMed PMID: 25890745; PMCID: PMC4405660.

14. Nelson DM, Hashizume R, Yoshizumi T, Blakney AK, Ma Z, Wagner WR. Intramyocardial injection of a synthetic hydrogel with delivery of bFGF and IGF1 in a rat model of ischemic cardiomyopathy. Biomacromolecules. 2014;15(1):1–11. Epub 20140102. doi: 10.1021/bm4010639. PubMed PMID: 24345287; PMCID: PMC4214385.

15. Freytes DO, Martin J, Velankar SS, Lee AS, Badylak SF. Preparation and rheological characterization of a gel form of the porcine urinary bladder matrix. Biomaterials. 2008;29(11):1630–7. Epub 20080116. doi: 10.1016/j.biomaterials.2007.12.014. PubMed PMID: 18201760.

16. Hashizume R, Hong Y, Takanari K, Fujimoto KL, Tobita K, Wagner WR. The effect of polymer degradation time on functional outcomes of temporary elastic patch support in ischemic cardiomyopathy. Biomaterials. 2013;34(30):7353–63. Epub 20130701. doi: 10.1016/j.biomaterials.2013.06.020. PubMed PMID: 23827185; PMCID: PMC3804157.

17. Carpentier A, Chachques JC. Myocardial substitution with a stimulated skeletal muscle: first successful clinical case. Lancet. 1985;1(8440):1267. doi: 10.1016/s0140-6736(85)92329-3. PubMed PMID: 2860458.

18. Batista RJ, Verde J, Nery P, Bocchino L, Takeshita N, Bhayana JN, Bergsland J, Graham S, Houck JP, Salerno TA. Partial left ventriculectomy to treat end-stage heart disease. Ann Thorac Surg. 1997;64(3):634–8. doi: 10.1016/s0003-4975(97)00779-0. PubMed PMID: 9307450.

19. Chaudhry PA, Mishima T, Sharov VG, Hawkins J, Alferness C, Paone G, Sabbah HN. Passive epicardial containment prevents ventricular remodeling in heart failure. Ann Thorac Surg. 2000;70(4):1275–80. doi: 10.1016/s0003-4975(00)01755-0. PubMed PMID: 11081885.

20. Hetzer R, Javier M, Wagner F, Loebe M, Javier Delmo EM. Organ-saving surgical alternatives to treatment of heart failure. Cardiovasc Diagn Ther. 2021;11(1):213–25. doi: 10.21037/cdt-20-285. PubMed PMID: 33708494; PMCID: PMC7944209.

21. Kittleson MM, Kobashigawa JA. Cardiac Transplantation: Current Outcomes and Contemporary Controversies. JACC Heart Fail. 2017;5(12):857–68. doi: 10.1016/j.jchf.2017.08.021. PubMed PMID: 29191293.

22. Ondrusek M, Artemiou P, Bezak B, Gasparovic I, By TM, Durdik S, Lesny P, Goncalvesova E, Hulman M. Temporal Analysis in Outcomes of Long-Term Mechanical Circulatory Support: Retrospective Study. Thorac Cardiovasc Surg. 2024. Epub 20240419. doi: 10.1055/s-0044-1782600. PubMed PMID: 38641334.

23. Broughton KM, Wang BJ, Firouzi F, Khalafalla F, Dimmeler S, Fernandez-Aviles F, Sussman MA. Mechanisms of Cardiac Repair and Regeneration. Circ Res. 2018;122(8):1151–63. doi: 10.1161/circresaha.117.312586. PubMed PMID: 29650632; PMCID: PMC6191043.

24. Michler RE. The current status of stem cell therapy in ischemic heart disease. J Card Surg. 2018;33(9):520–31. Epub 20180822. doi: 10.1111/jocs.13789. PubMed PMID: 30136308.

25. Huang K, Hu S, Cheng K. A New Era of Cardiac Cell Therapy: Opportunities and Challenges. Adv Healthc Mater. 2019;8(2):e1801011. Epub 20181213. doi: 10.1002/adhm.201801011. PubMed PMID: 30548836; PMCID: PMC6368830.

26. Jackson BM, Gorman JH, 3rd, Salgo IS, Moainie SL, Plappert T, St John-Sutton M, Edmunds LH, Jr., Gorman RC. Border zone geometry increases wall stress after myocardial infarction: contrast echocardiographic assessment. Am J Physiol Heart Circ Physiol. 2003;284(2):H475–9. Epub 20021031. doi: 10.1152/ajpheart.00360.2002. PubMed PMID: 12414441.

27. Chen QZ, Bismarck A, Hansen U, Junaid S, Tran MQ, Harding SE, Ali NN, Boccaccini AR. Characterisation of a soft elastomer poly(glycerol sebacate) designed to match the mechanical properties of myocardial tissue. Biomaterials. 2008;29(1):47–57. doi: 10.1016/j.biomaterials.2007.09.010. PubMed PMID: 17915309.

28. Lee LC, Wall ST, Klepach D, Ge L, Zhang Z, Lee RJ, Hinson A, Gorman JH, 3rd, Gorman RC, Guccione JM. Algisyl-LVR™ with coronary artery bypass grafting reduces left ventricular wall stress and improves function in the failing human heart. Int J Cardiol. 2013;168(3):2022–8. Epub 20130208. doi: 10.1016/j.ijcard.2013.01.003. PubMed PMID: 23394895; PMCID: PMC3748222.

29. Johnson TD, Lin SY, Christman KL. Tailoring material properties of a nanofibrous extracellular matrix derived hydrogel. Nanotechnology. 2011;22(49):494015. Epub 20111121. doi: 10.1088/0957-4484/22/49/494015. PubMed PMID: 22101810; PMCID: PMC3280097.

30. U. Zeymer, S.V. Rao, M.W. Krucoff, A Placebo-controlled, Multicenter, Randomized, Doubleblind Trial to Evaluate the Safety and Effectiveness of IK-5001 (Bioabsorbable Cardiac Matrix [BCM]) for the Prevention of Remodeling of the Ventricle and Congestive Heart Failure after Acute Myocardial Infarction, 2015. https://www.escardio.org/static_!le/Escardio/Press-media/Press%20releases/2015/Congress/PRESERVATION_Zeymer.pdf (Accessed 30 October 2016).

31. Mann DL, Lee RJ, Coats AJ, Neagoe G, Dragomir D, Pusineri E, Piredda M, Bettari L, Kirwan BA, Dowling R, Volterrani M, Solomon SD, Sabbah HN, Hinson A, Anker SD. One-year follow-up results from AUGMENT-HF: a multicentre randomized controlled clinical trial of the efficacy of left ventricular augmentation with Algisyl in the treatment of heart failure. Eur J Heart Fail. 2016;18(3):314–25. Epub 20151111. doi: 10.1002/ejhf.449. PubMed PMID: 26555602.

32. Traverse JH, Henry TD, Dib N, Patel AN, Pepine C, Schaer GL, DeQuach JA, Kinsey AM, Chamberlin P, Christman KL. First-in-Man Study of a Cardiac Extracellular Matrix Hydrogel in Early and Late Myocardial Infarction Patients. JACC Basic Transl Sci. 2019;4(6):659–69. Epub 20190911. doi: 10.1016/j.jacbts.2019.07.012. PubMed PMID: 31709316; PMCID: PMC6834965.

33. Wall ST, Walker JC, Healy KE, Ratcliffe MB, Guccione JM. Theoretical impact of the injection of material into the myocardium: a finite element model simulation. Circulation. 2006;114(24):2627–35. Epub 20061127. doi: 10.1161/circulationaha.106.657270. PubMed PMID: 17130342.

34. Li DS, Avazmohammadi R, Rodell CB, Hsu EW, Burdick JA, Gorman JH, 3rd, Gorman RC, Sacks MS. How hydrogel inclusions modulate the local mechanical response in early and fully formed post-infarcted myocardium. Acta Biomater. 2020;114:296–306. Epub 20200730. doi: 10.1016/j.actbio.2020.07.046. PubMed PMID: 32739434; PMCID: PMC7484038.

35. Ifkovits JL, Tous E, Minakawa M, Morita M, Robb JD, Koomalsingh KJ, Gorman JH, 3rd, Gorman RC, Burdick JA. Injectable hydrogel properties influence infarct expansion and extent of postinfarction left ventricular remodeling in an ovine model. Proc Natl Acad Sci U S A. 2010;107(25):11507–12. Epub 20100607. doi: 10.1073/pnas.1004097107. PubMed PMID: 20534527; PMCID: PMC2895138.

36. Rodell CB, Lee ME, Wang H, Takebayashi S, Takayama T, Kawamura T, Arkles JS, Dusaj NN, Dorsey SM, Witschey WR, Pilla JJ, Gorman JH, 3rd, Wenk JF, Burdick JA, Gorman RC. Injectable Shear-Thinning Hydrogels for Minimally Invasive Delivery to Infarcted Myocardium to Limit Left Ventricular Remodeling. Circ Cardiovasc Interv. 2016;9(10). doi: 10.1161/circinterventions.116.004058. PubMed PMID: 27729419; PMCID: PMC5123705.

37. Plotkin M, Vaibavi SR, Rufaihah AJ, Nithya V, Wang J, Shachaf Y, Kofidis T, Seliktar D. The effect of matrix stiffness of injectable hydrogels on the preservation of cardiac function after a heart attack. Biomaterials. 2014;35(5):1429–38. Epub 20131121. doi: 10.1016/j.biomaterials.2013.10.058. PubMed PMID: 24268664.

38. Wang H, Rodell CB, Lee ME, Dusaj NN, Gorman JH, 3rd, Burdick JA, Gorman RC, Wenk JF. Computational sensitivity investigation of hydrogel injection characteristics for myocardial support. J Biomech. 2017;64:231–5. Epub 20170901. doi: 10.1016/j.jbiomech.2017.08.024. PubMed PMID: 28888476; PMCID: PMC5694362.

39. Griffin DR, Weaver WM, Scumpia PO, Di Carlo D, Segura T. Accelerated wound healing by injectable microporous gel scaffolds assembled from annealed building blocks. Nat Mater. 2015;14(7):737–44. Epub 20150601. doi: 10.1038/nmat4294. PubMed PMID: 26030305; PMCID: PMC4615579.

40. Sussman EM, Halpin MC, Muster J, Moon RT, Ratner BD. Porous implants modulate healing and induce shifts in local macrophage polarization in the foreign body reaction. Ann Biomed Eng. 2014;42(7):1508–16. Epub 20131119. doi: 10.1007/s10439-013-0933-0. PubMed PMID: 24248559.

41. Koch B, Sanchez S, Schmidt CK, Swiersy A, Jackson SP, Schmidt OG. Confinement and deformation of single cells and their nuclei inside size-adapted microtubes. Adv Healthc Mater. 2014;3(11):1753–8. Epub 20140425. doi: 10.1002/adhm.201300678. PubMed PMID: 24764273; PMCID: PMC4227890.

42. Han Y, Lian M, Wu Q, Qiao Z, Sun B, Dai K. Effect of Pore Size on Cell Behavior Using Melt Electrowritten Scaffolds. Front Bioeng Biotechnol. 2021;9:629270. Epub 20210702. doi: 10.3389/fbioe.2021.629270. PubMed PMID: 34277578; PMCID: PMC8283809.

43. Béduer A, Braschler T, Peric O, Fantner GE, Mosser S, Fraering PC, Benchérif S, Mooney DJ, Renaud P. A compressible scaffold for minimally invasive delivery of large intact neuronal networks. Adv Healthc Mater. 2015;4(2):301–12. Epub 20140901. doi: 10.1002/adhm.201400250. PubMed PMID: 25178838.

44. He T, Li B, Colombani T, Joshi-Navare K, Mehta S, Kisiday J, Bencherif SA, Bajpayee AG. Hyaluronic Acid-Based Shape-Memory Cryogel Scaffolds for Focal Cartilage Defect Repair. Tissue Eng Part A. 2021;27(11-12):748–60. Epub 20210205. doi: 10.1089/ten.TEA.2020.0264. PubMed PMID: 33108972.

45. Dobaczewski M, Gonzalez-Quesada C, Frangogiannis NG. The extracellular matrix as a modulator of the inflammatory and reparative response following myocardial infarction. J Mol Cell Cardiol. 2010;48(3):504–11. Epub 20090723. doi: 10.1016/j.yjmcc.2009.07.015. PubMed PMID: 19631653; PMCID: PMC2824059.

46. Zhao ZQ, Puskas JD, Xu D, Wang NP, Mosunjac M, Guyton RA, Vinten-Johansen J, Matheny R. Improvement in cardiac function with small intestine extracellular matrix is associated with recruitment of C-kit cells, myofibroblasts, and macrophages after myocardial infarction. J Am Coll Cardiol. 2010;55(12):1250–61. doi: 10.1016/j.jacc.2009.10.049. PubMed PMID: 20298933.

47. Nahrendorf M, Swirski FK. Monocyte and macrophage heterogeneity in the heart. Circ Res. 2013;112(12):1624–33. doi: 10.1161/circresaha.113.300890. PubMed PMID: 23743228; PMCID: PMC3753681.

48. Seif-Naraghi SB, Singelyn JM, Salvatore MA, Osborn KG, Wang JJ, Sampat U, Kwan OL, Strachan GM, Wong J, Schup-Magoffin PJ, Braden RL, Bartels K, DeQuach JA, Preul M, Kinsey AM, DeMaria AN, Dib N, Christman KL. Safety and efficacy of an injectable extracellular matrix hydrogel for treating myocardial infarction. Sci Transl Med. 2013;5(173):173ra25. doi: 10.1126/scitranslmed.3005503. PubMed PMID: 23427245; PMCID: PMC3848875.

49. Sack KL, Aliotta E, Choy JS, Ennis DB, Davies NH, Franz T, Kassab GS, Guccione JM. Intra-myocardial alginate hydrogel injection acts as a left ventricular mid-wall constraint in swine. Acta Biomater. 2020;111:170–80. Epub 20200516. doi: 10.1016/j.actbio.2020.04.033. PubMed PMID: 32428678; PMCID: PMC7368390.

50. Wang H, Rodell CB, Zhang X, Dusaj NN, Gorman JH, 3rd, Pilla JJ, Jackson BM, Burdick JA, Gorman RC, Wenk JF. Effects of hydrogel injection on borderzone contractility post-myocardial infarction. Biomech Model Mechanobiol. 2018;17(5):1533–42. Epub 20180531. doi: 10.1007/s10237-018-1039-2. PubMed PMID: 29855734.

